# Mapping the landscape of transcription factor promoter activity during vegetative development in Marchantia

**DOI:** 10.1101/2023.06.17.545419

**Authors:** Facundo Romani, Susanna Sauret-Güeto, Marius Rebmann, Davide Annese, Ignacy Bonter, Marta Tomaselli, Tom Dierschke, Mihails Delmans, Eftychios Frangedakis, Linda Silvestri, Jenna Rever, John L. Bowman, Ignacio Romani, Jim Haseloff

## Abstract

Transcription factors (TFs) are essential for the regulation of gene expression and cell fate determination. Characterising the transcriptional activity of TF genes in space and time is a critical step towards understanding complex biological systems. The vegetative gametophyte meristems of bryophytes share some characteristics with the shoot-apical meristems of flowering plants. However, the identity and expression profiles of TFs associated with gametophyte organization are largely unknown. With only ∼450 TF genes, *Marchantia polymorpha* is an outstanding model system for plant systems biology. We have generated a near-complete collection of promoter elements derived from Marchantia TF genes. We experimentally tested *in planta* reporter fusions for all the TF promoters in the collection and systematically analysed expression patterns in Marchantia gemmae. This allowed us to build a map of precise expression domains and identify a unique set of TFs expressed in the stem-cell zone, providing new insight into the dynamic regulation of the gametophytic meristem and its evolution. In addition, we provide an online database of expression patterns for all promoters in the collection. We expect that the promoter elements characterised here will be useful for cell-type specific expression, synthetic biology applications, and functional genomics.

## INTRODUCTION

Embryophytes evolved around 470 million years ago and started covering the Earth’s land surface. A common feature of the body plan of land plants is the alternation of generations between the sporophyte and the gametophyte during vegetative to reproductive development (Bowman et al., 2016; Bowman, 2022b). The major lineages display two contrasting forms of vegetative body: tracheophytes (vascular plants) display a dominant sporophyte generation (diploid), while the vegetative body of bryophytes is gametophytic (haploid). Both tissues are characterised by polar growth with apical dominance and maintenance of a stem-cell population. Developmental programs controlling meristem organization in the sporophyte of vascular plants are relatively well known (Lodha et al., 2008; Uchida and Torii, 2019). It is expected that the vegetative body of bryophytes has an equivalent meristem organization, but the regulatory programs associated with the bryophyte gametophyte and how it evolved are not fully understood (Bowman et al., 2019; Hata and Kyozuka, 2021).

During the last decade, evo-devo studies in models such as *Marchantia polymorpha* and *Physcomitrium patens* have provided exceptional insights into the molecular mechanisms regulating developmental programs in bryophytes. Several aspects of hormonal and peptide signalling follow strikingly similar rules to flowering plants (Blazquez et al., 2020; Hirakawa, 2022). However, less is known about the identity of transcription factors (TFs) regulating vegetative development of the apical meristem of bryophytes (Romani and Moreno, 2021). A better understanding of the nature of these two forms of vegetative growth is likely to shed light on the early evolution of land plants.

TFs are key determinants of genetic programs operating during cellular development, and their cell-type specific patterns of expression provide indicators for regulatory processes that underpin cell differentiation during the vegetative body formation. *M. polymorpha* shows many advantages as an experimental system and has become a significant model organism for plant science (Kohchi et al., 2021; Bowman, 2022a; Bowman et al., 2022). Marchantia not only widens our knowledge of plant molecular biology outside of flowering plants (angiosperms) but is also an exceptional model for synthetic biology (Boehm et al., 2017; Sauret-Gueto et al., 2020). The Marchantia genome features only about ∼450 TF genes (Bowman et al., 2017), about a fifth of the number of TFs in *A. thaliana*, with several subfamilies containing a single gene. This greatly simplifies the study of complex gene regulatory networks (GRN), which is afflicted by the problem of gene redundancy in other systems (Wagner, 1996; Panchy et al., 2016). Combined with a short haploid life cycle and efficient *Agrobacterium*-mediated transformation protocols (Ishizaki et al., 2016), fast modular growth, and simple morphology (Boehm et al., 2017), Marchantia allows system-wide experimental approaches that are infeasible in other plant species.

The mapping of temporal and spatial gene expression patterns is essential for understanding regulatory networks underlying developmental processes. In the last few years, different techniques have been developed to explore gene expression using single-cell (scRNA-seq) and spatial transcriptomics (Giacomello, 2021; Seyfferth et al., 2021; Wang et al., 2023). These techniques can provide gene expression information at a single-cell level for an entire transcriptome but associating that to cell identities present some challenges and limitations (Yuan et al., 2017). On the other hand, traditional tools, such as using transgenic lines with reporters fused to predicted promoter regions, can deliver a more detailed map of expression patterns at the cellular level. This approach can provide insight into the dynamics of gene expression as well as useful tools for tissue-specific expression. However, understanding the landscape of gene expression in an organism requires exploring the expression of hundreds to thousands of genes. The generation of stable transgenic lines is laborious and time-consuming, making such an endeavour infeasible for many model organisms, particularly in plant species. Yet, comprehensive expression pattern databases have been established for several metazoan species using transcriptional reporters and *in situ* hybridisation (Visel et al., 2007; Gallo et al., 2011; Bessa et al., 2014; Alonso-Barba et al., 2016).

In this work, we aimed to systematically explore the behaviour of promoter elements from TF genes in Marchantia, and to map the resulting expression patterns. We hypothesise the gametophytic meristem is also characterised by the specific expression patterns of TFs in Marchantia and they could provide clues to understand underlying GRNs. We characterised a near-complete collection of promoter elements derived from TFs encoded in the genome of Marchantia. These patterns were used as surrogates for the underlying gene circuits and enabled us to survey the regulatory landscape in the vegetative gametophytes of Marchantia. The approach offers an unbiased way to explore TF expression patterns in the meristem. Comparative analysis of the reporters allowed us to recognise expression domains and cell types in Marchantia gemmae and provide important insights into the genetic programs underpinning the organization of Marchantia stem cells. We also identified cell-type specific promoters across different stages of gemma development. Surprisingly, the set of TF reporters found in in the stem-cell zone is largely evolutionary unrelated to TFs known from the sporophyte meristem of vascular plants. The imaging data for all tested promoters is available via a web-based database to accelerate functional genomics studies and cell-type specific engineering.

## RESULTS

### A comprehensive collection of putative promoters from Marchantia TF genes

We built a library of synthetic promoters derived from the upstream region of TF genes identified in the *M. polymorpha* Tak-1 v5.1 genome (Bowman et al., 2017; Montgomery et al., 2020). For each TF we synthesised the 5’UTR region (5UTR) and putative promoter regions (PROM) of ∼1.8 kb from the annotated transcription start site (TSS). 5UTRs shorter than 500 bp were cloned as a single unit (PROM5), while longer than 500 bp and smaller than 3kb, were synthesised as separate 5UTR and PROM parts (Fig. 1b). In average, the length of the promoters cloned is about 2.5kb (Fig. 1a). The collection is widely distributed across all major plant TF protein families (Fig. 1c), with a total coverage of around ∼82% of all TFs in the Marchantia genome. In addition, the collection includes promoters for other relevant genes in Marchantia that serve as references (Supplemental Table S1). Promoter sequences were domesticated following the standards for Type IIS cloning and inserted as L0 parts for Loop assembly (Patron et al., 2015; Pollak et al., 2019) to facilitate the reuse of the synthetic parts.

**Figure 1.**
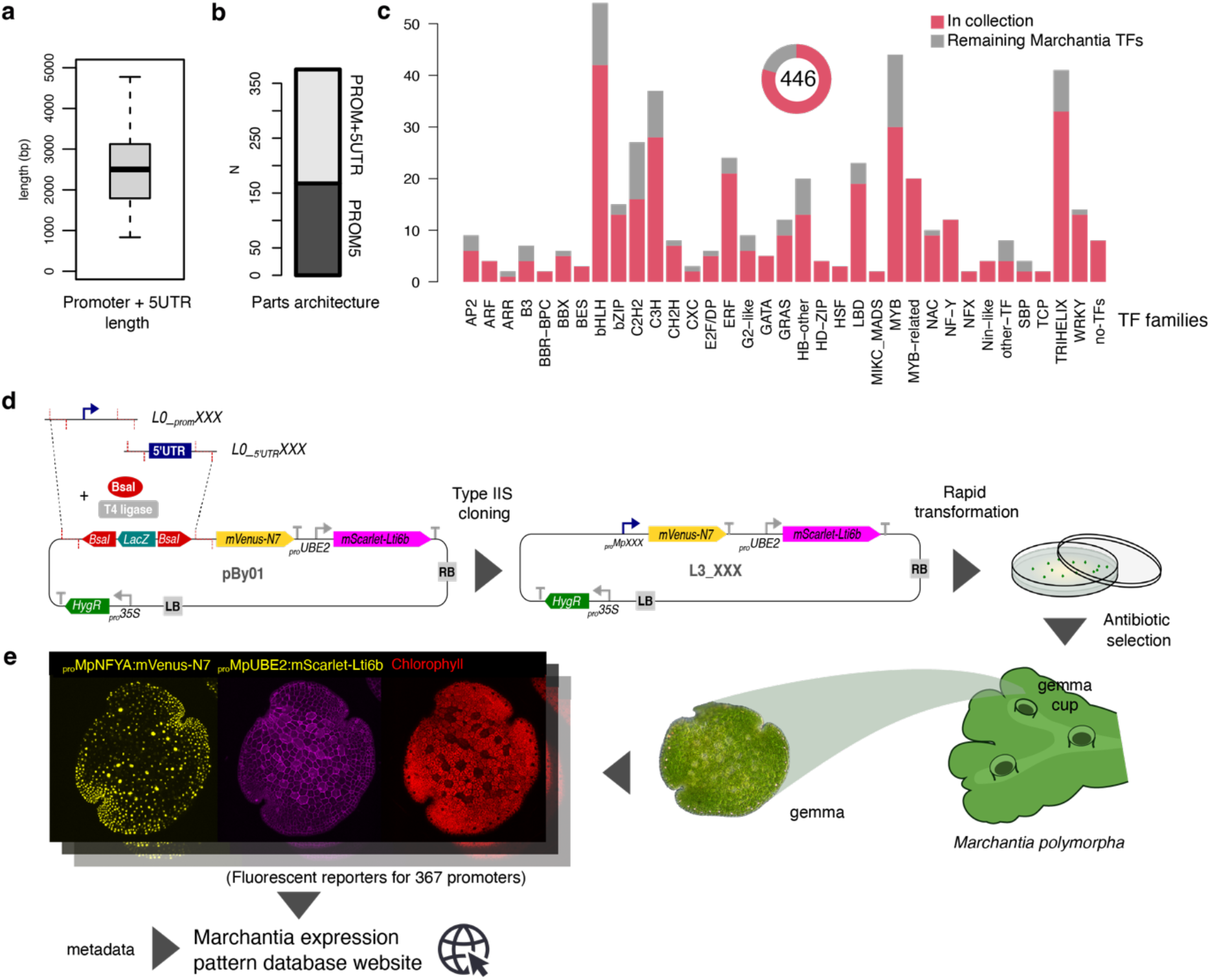
Overview of the transcription factor promoter collection. (a) Boxplot showing the length distribution of all promoters in the collection and (b) the architecture of the synthetic parts. (c) Distribution of tested TFs (red) across TF families in the Marchantia genome. (d) Overview of the cloning and transformation strategy implemented to characterize the promoters, including (e) an example (*_pro_*Mp*NFYA, Mp1g13740*) of the imaging output for each promoter deposited in an accessible database.

We subsequentially cloned the promoter elements into a binary vector containing an mVenus fluorescent protein with an N7 nuclear localization signal to drive the expression of the promoter of interest and a plasma membrane marker (mScarlet-Lti6b) controlled by the *_pro_MpUBE2* constitutive promoter (Sauret-Gueto et al., 2020) as part of the same T-DNA insertion cassette (Fig. 1d). This marker works as a positive control for the transformation, an internal reference for any artifacts associated with the insertion site of the construct and helped to visualize different cell shapes and arrangements and to classify patterns. To avoid the intermediate cloning steps, we built a custom vector with Type IIS sites for cloning of PROM5 or PROM and 5UTR L0 parts in a backbone with pre-assembled parts, obtaining the desired final construct for stable expression in a single step (Fig. 1d). Finally, we implemented a high-throughput transformation protocol based on *Agrobacterium*-mediated transformation in multi-well plates (Ishizaki et al., 2008; Sauret-Gueto et al., 2020) to obtain 6−7 independent stable transgenic lines for each plasmid.

### Characterising TF reporters in planta

Marchantia produces vegetative propagules called gemmae, which have a lenticular disc-like morphology and accumulate in cups. Gemmae provide a stereotypically conserved initial stage of the Marchantia vegetative life cycle, with typically two opposing apical notches containing stem-cells, differentiated cells, two planes of symmetry and no pre-defined abaxial/adaxial polarity (Kato et al., 2020a; Zheng et al., 2020). During the gemma stage, the structure of the stem-cell niche and the entire body is accessible for microscopy and differentiating cells can be recognised easily without the need for staining or clearing.

We imaged several lines for each promoter using confocal microscopy and selected images that best represent the consensus expression patterns. From the collection of 367 promoters, we initially classified the expression patterns into 5 non-exclusive categories: 104 lines showed no detectable signal (28%), 41 presented a dim signal (11%), 121 a ubiquitous expression pattern across the gemma (33%), 127 a pattern stronger or specific to the notch area (35%), and, finally, 50 (13%) had some specificity for specialised cells (Supplemental Table S1).

To test whether these expression patterns correlate with endogenous expression, we compared each group with transcript levels from the corresponding genes analysed by RNA-seq analysis in whole gemmae (Mizuno et al., 2021). As expected, ubiquitous promoters showed the highest average TPM values, followed by genes associated with specialised cells and notch biased expression (Fig. 2A-B). On the other hand, reporters with no expression had the lowest TPM values, followed by the group with dim expression (poor signal-to-noise ratio). From this latter group of TFs, several genes had higher expression levels in other developmental stages (Kawamura et al., 2022). Only around ∼15% presented clearly inconsistent expression patterns compared to RNA-seq.

**Figure 2.**
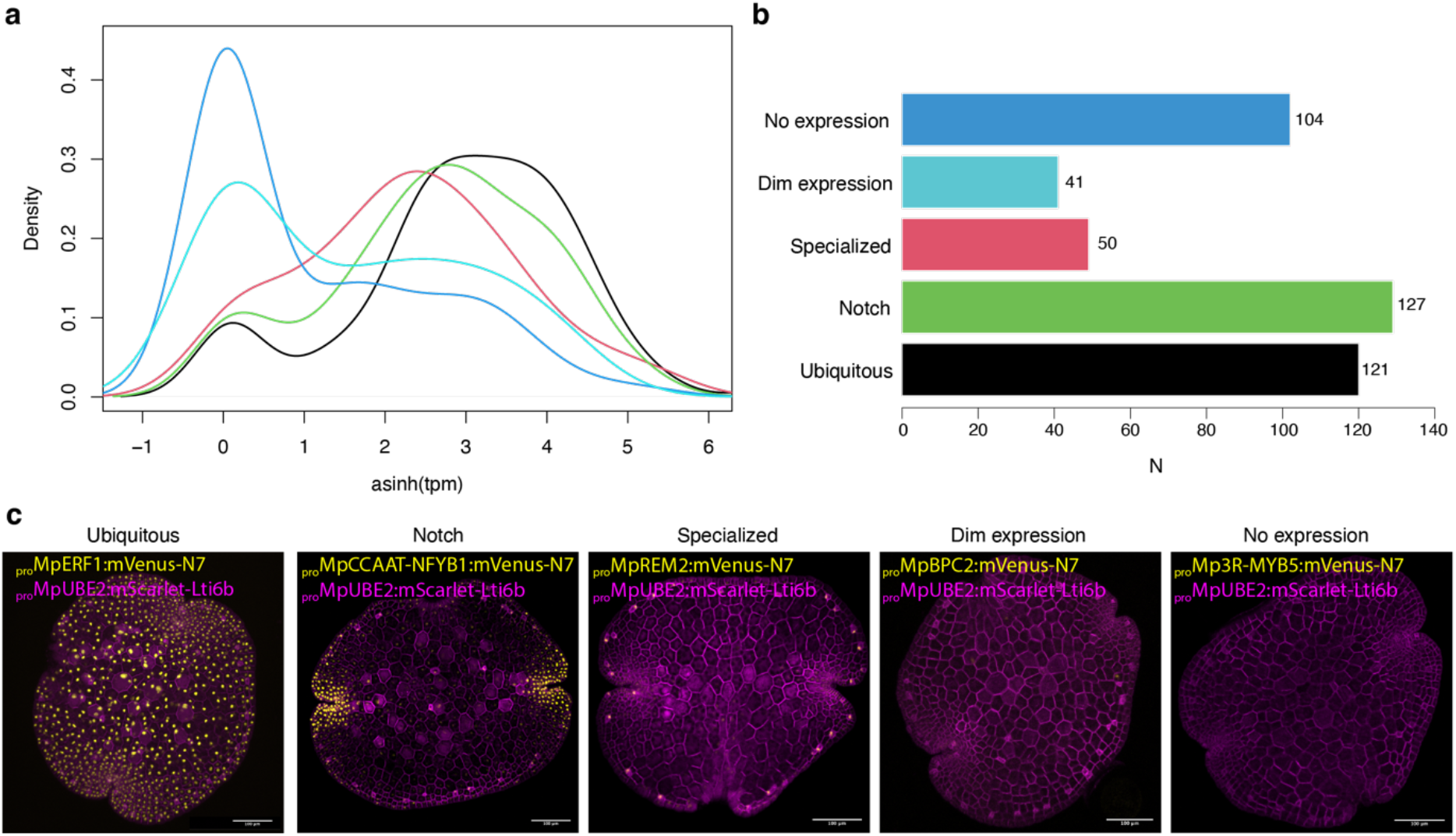
Quality control for the promoter collection. (a) Density plot of initial TF classifications: no expression, dim expression, specialised, notch, and ubiquitous across asinh(TPM) values from whole tissue RNA-seq of the gemma. (b) Number of unique promoters tested in each class. (c) Examples of promoters belonging to each class (*_pro_*Mp*ERF1, _pro_*Mp*CCAAT-NFYB1, _pro_*Mp*REM2, _pro_*Mp*BPC2, _pro_*Mp*3R-MYB5*). Confocal images of the gene of interest (yellow) and a constitutive plasma membrane marker (magenta, *_pro_*Mp*UBE2:mScarlet-Lti6b*). Scale bar 100 μm. Gene IDs: Mp*ERF1 = Mp1g20040, _pro_*Mp*CCAAT-NFYB1 = Mp4g13360, _pro_*Mp*REM2 = Mp2g08790, _pro_*Mp*BPC2 = MpVg00350, _pro_*Mp*3R-MYB5 = Mp4g04750*.

The microscopy data collected during the screening of promoter activities have been organised in a database accessible online (Fig. 1, https://mpexpatdb.org/). The collection can be searched and filtered by expression profiles, gene IDs, names, and families. The database links promoters with functional information about the adjacent gene available in the MarpolBase (Ishizaki et al., 2016). For each reporter construct tested we recorded a maximum projection image with 3 separate channels (gene of interest, chlorophyll autofluorescence, and the constitutive plasma membrane marker) for identification of cell types. We have also developed an original feature to visualize the channels independently. The user can select which channels are actively visualised and download the appropriate composite picture.

### Identifying expression domains in Marchantia gemmae

The variability between individuals is relatively low and the dimensions of the tissue follow a normal distribution (Fig. S1). This simple morphology makes the gemma stage convenient for systematic comparisons between reporters. Excluding promoters with dim or no expression levels, for each representative reporter we orientated the image to align the two apical notches to the horizontal axis, subtracted the background, and made a profile of the fluorescence intensity along the notch axis. The length of the profile was adjusted to fit the notches at the same distance and then smoothed to reduce the noise of the signal. To avoid small variations between left and right notch, we averaged them. Finally, we normalised the signal to the maximum of each image (Fig 3a). This allowed us to generate a linear vector that represented expression patterns from different transgenic lines in a comparable way.

**Figure 3.**
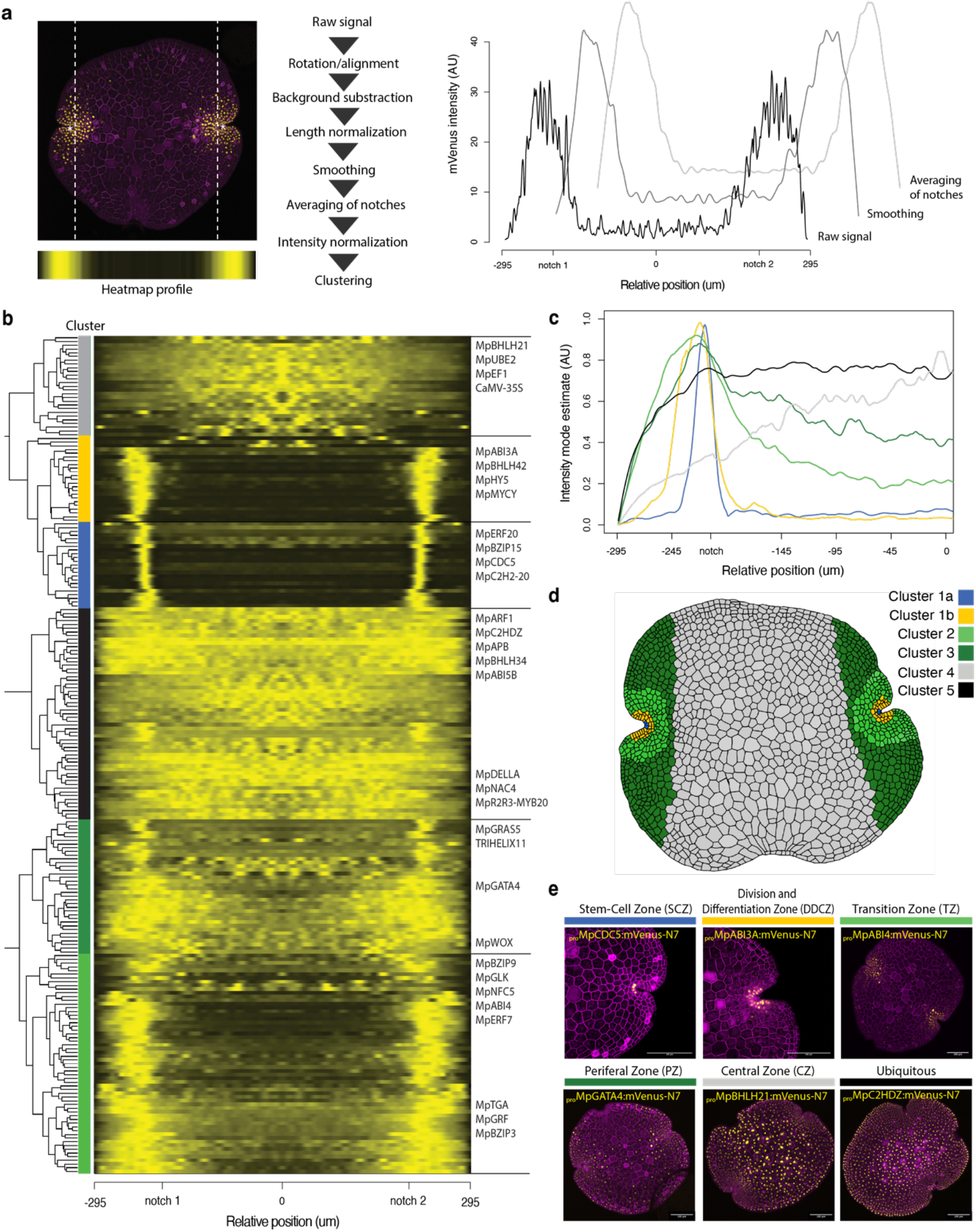
Clustering analysis of expression patterns in Marchantia gemmae. (a) Pipelines for image processing of confocal images to obtain normalised profiles to compare expression patterns between gemmae (see Methods section for detail). Example of confocal images of a fluorescent reporter (left), the corresponding plot of profiles for intermediate steps of the pipeline (right) and heatmap (bottom left). (b) Heatmap of promoters with significant expression and dendrogram of hierarchical clustering with the following color code: Blue, cluster 1a; Yellow, cluster 1b; Light green, cluster 2; Dark green, cluster 3; Grey, cluster 4; Black, cluster 5. (c) Mode of the profile for each cluster across the gemmae. (d) Schematic map of the association of each cluster with distinct cellular expression domains in the Marchantia gemma. (e) Example of TF fluorescent reporters for each cluster (*_pro_*Mp*CDC5, _pro_*Mp*ABI3A, _pro_*Mp*ABI4, _pro_*Mp*GATA4, _pro_*Mp*BHLH21, _pro_*Mp*C2HDZ*). Confocal images of the gene of interest (yellow) and a constitutive plasma membrane marker (magenta, *_pro_*Mp*UBE2:mScarlet-Lti6b*). Scale bar 100 μm. Gene IDs: Mp*CDC5 = Mp1g10310,* Mp*ABI3A = Mp5g08310,* Mp*ABI4 = Mp7g00860,* Mp*GATA4 = Mp7g03490,* Mp*BHLH21 = Mp3g11900,* Mp*C2HDZ = Mp2g24200*.

In total, we analysed reporters for 218 different genes. We used hierarchical clustering and identified 5 clusters representing distinct expression domains (Fig. 3b-e). Most expression patterns follow a skewed distribution with the apical notch position as the mode (cluster 1-3). Others instead followed a normal distribution with the central zone as the mode (cluster 4) or were evenly distributed across the gemma (cluster 5). Only a few expression patterns did not match these broad classes, and these were mostly associated with expression in differentiated scattered cells.

Within cluster 1, we distinguished two populations, one with a peak in the apical notch and a second includes a broader area around it (Fig. 3c). These correspond to the stem-cell zone (SCZ) and Dividing and Differentiating Cell Zone (DDCZ) respectively, as recognised earlier (Kohchi et al., 2021). The SCZ includes a single apical cell and sub-apical cell anticlinal derivatives located at the center of the notch (Kohchi et al., 2021). The DDCZ covers a population of two rows of derivative cells precisely arranged around the SCZ. Cluster 2 is a broader area of small cells radially distributed along the SCZ that we named Transition Zone (TZ). Cluster 3 also includes the previous domains but extends over a group of cells distant to the apical notches and fades along the axis. We named this domain of larger cells peripheral zone (PZ). Finally, we named Cluster 4 and 5 that correspond to two populations of ubiquitous promoters with different strengths between the apical region and the central zone (CZ). Most known constitutive promoters (*_pro_*Mp*UBE2, _pro_*Mp*EF1, _pro_CaMV35S*) belong to cluster 4 (Althoff et al., 2014; Sauret-Gueto et al., 2020). Finally, based on clustering analysis and incorporating literature information about cell-types in Marchantia, we generated a schematic model of a gemma that described cellular arrangements and cell populations that could be distinguished (Fig. 3d).

We selected reporters representative of expression domains and cell-types to obtain a more precise map of the expression domains at a cellular level. We built transgenic lines with different combinations of promoters driving the expression of two or three compatible fluorescent reporters (mVenus, mScarlet and mTurquoise) localised in the nucleus as part of the same T-DNA. In all cases, the domains could be clearly distinguished in the different combinations (Fig. 4). This demonstrates that the expression patterns could be used in an independent and additive fashion to mark multiple cell states simultaneously and allowed us to differentiate between promoters active in the SCZ and the apical cell in the middle (Fig. 4a,d).

**Figure 4.**
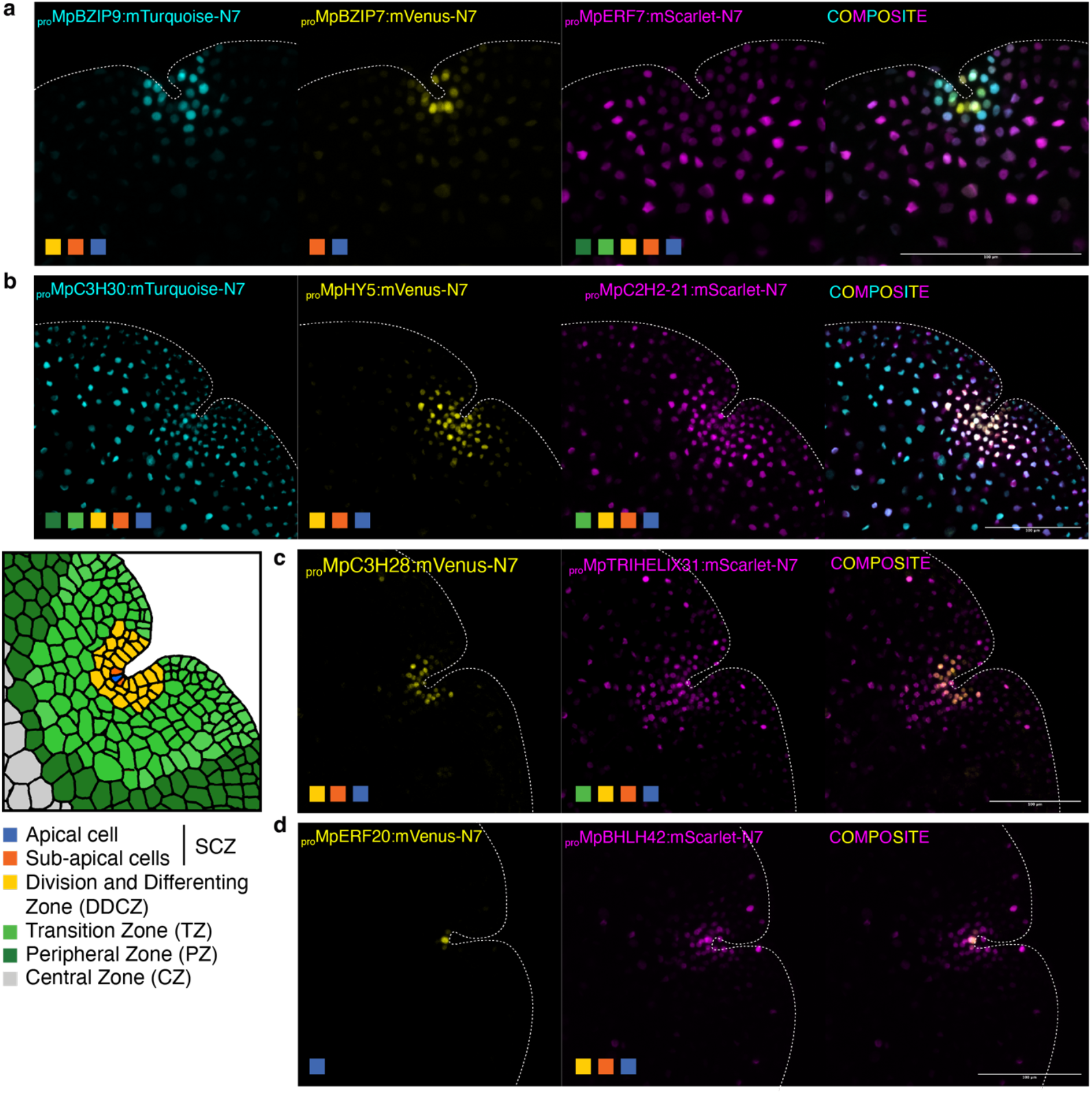
Combination of multiple fluorescent reporters. Confocal images of the apical region of Marchantia gemmae transformed with multiple fluorescent reporters of TFs in the same plasmid. (a) Combo 1: *_pro_*Mp*BZIP9, _pro_*Mp*BZIP7, _pro_*Mp*ERF7*. (b) Combo 2: *_pro_*Mp*C3H30, _pro_*Mp*HY5, _pro_*Mp*C2H2-21*. (c) Combo 3: *_pro_*Mp*C3H28, _pro_*Mp*TRIHELIX31*. (d) Combo 4: *_pro_*Mp*ERF20, _pro_*Mp*BHLH42*. Schematic map and legend of the expression domains and cell-types in the gemma notch is shown (bottom left). Colour squares indicate the domains where each selected promoter is active. Individual channels and composite images are shown. Scale bar 100 μm. Gene IDs: Mp*BZIP9 = Mp6g03920,* Mp*BZIP7 = Mp3g04360,* Mp*ERF7 = Mp6g04880,* Mp*C3H30 = Mp7g18530,* Mp*HY5 = Mp1g16800,* Mp*C2H2-21 = Mp3g11570,* Mp*C3H28 = Mp7g14310,* Mp*TRIHELIX31 = Mp4g09730,* Mp*ERF20/LAXR =* Mp5g06970, Mp*BHLH42 = Mp5g09710*.

### Mapping TF expression patterns in specific cell-types

The global analysis of expression profiles along the apical axis can provide a systematic account of organism-wide patterns but may not capture the local cell patterning important for cell differentiation. To get a more precise map of cell types, we manually inspected each reporter and identified promoters with specificity for specialised cells such as rhizoids, oil body cells, and mucilage papillae (Figure 4b). In addition, two other expression domains (border and attachment) do not form regular distributions along the apical axis as most of the other domains (see below). These cell types and domains match descriptions in the published literature on cellular analysis in the Marchantia gemmae (Shimamura, 2016) and can be included in the schematic model gemma (Fig. 5a). The classification of cell types was defined in a way that any observed expression pattern could be classified as active in one or a combination of cell types. The corresponding TF gene families associated with the expression patterns are distributed across different cell-types (Fig. 5c). We did not find a clear association of a particular TF family with specific cell-types. Finally, clustering analysis of the expression domains and cell types reconstruct cell differentiation dynamics (Fig 5d).

**Figure 5.**
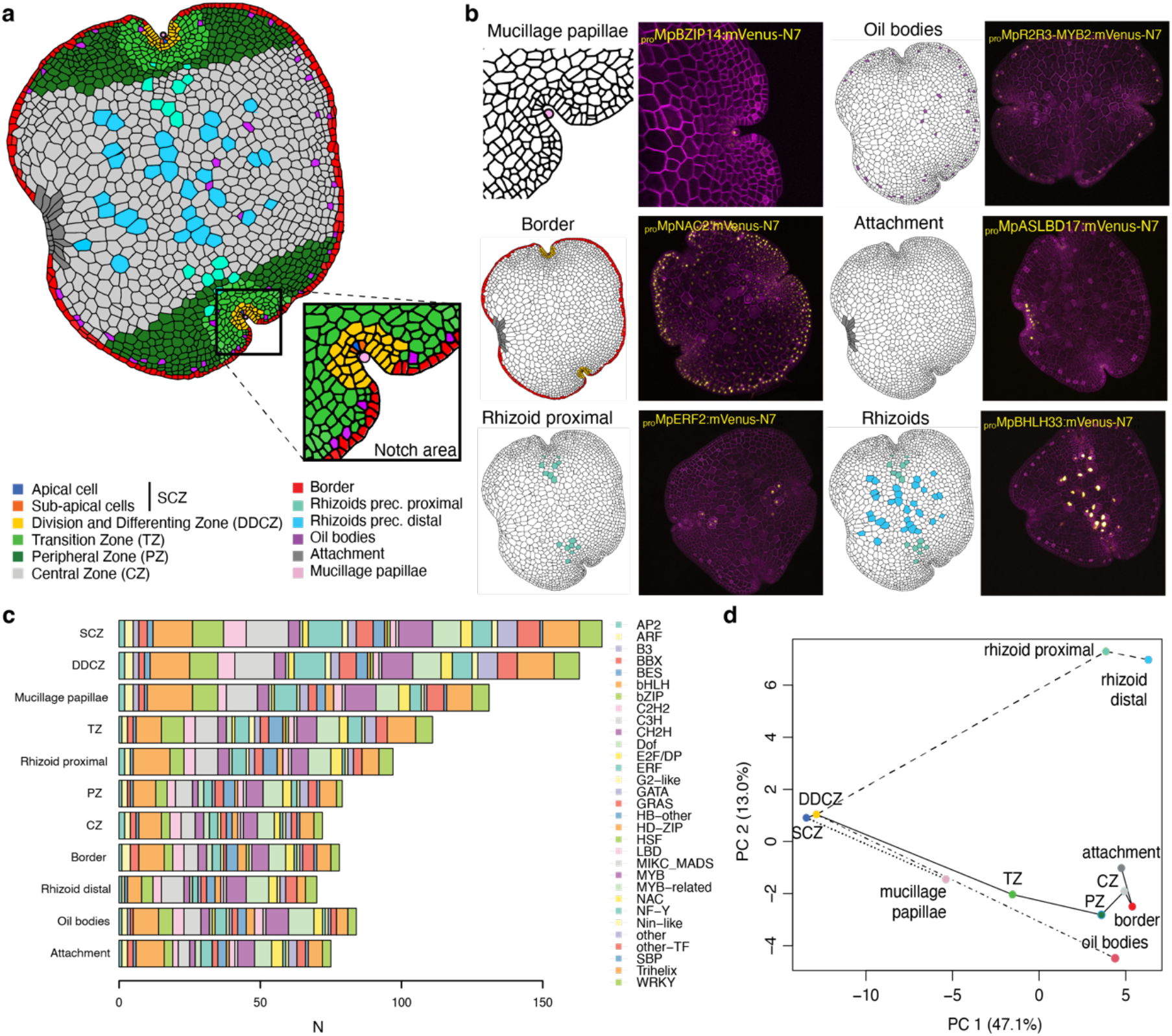
A model for promoter activity in the Marchantia gemmae. (a) Schematic representation of cell-types identified in the Marchantia gemma and (b) detailed view of the notch area. Examples of representative fluorescent reporters displaying cell-type specific expression patterns (B): *_pro_*Mp*BZIP14*, *_pro_*Mp*R2R3-MYB2*, *_pro_*Mp*NAC2 _pro_*Mp*ASLBD17*, *_pro_*Mp*ERF2*, and *_pro_*Mp*BHLH33/*Mp*RSL3*. Marked cell-types are shown (left) with confocal images (right) of the gene of interest (yellow) and a constitutive plasma membrane marker (magenta, *_pro_*Mp*UBE2:mScarlet-Lti6b*). (c) Number of reporters with expression across cell-types colored by TF gene families. (d) Principal component analysis (PCA) of cell-types based on the expression of TF reporters. Gene IDs: Mp*BZIP14 = Mp2g02230,* Mp*R2R3-MYB2 = Mp3g07510,* Mp*NAC2 = Mp6g02590,* Mp*ASLBD17 = Mp8g09250,* Mp*ERF2 = Mp7g13760,* Mp*BHLH33/*Mp*RSL3 = Mp1g01110*.

We identified promoters specific for cell lineages of specialised cells in Marchantia gemmae (Fig. 5b). Mucilage papillae are tip-growing cells covering the SCZ (Galatis and Apostolakos, 1977). We showed that *_pro_*Mp*BZIP14* and *_pro_*Mp*BHLH28* were specifically active in the mucilage papillae (Fig. 5b, Suppl. Table S1). Oil body cells are idiotypic cells scattered across the thallus and are distributed in regular fashion along the edges of gemmae (Romani et al., 2022). Our screening also led to the rediscovery of oil body-specific promoters for the genes Mp*ERF13*, Mp*C1HDZ*, and Mp*R2R3-MYB2* (Fig. 5b, Suppl. Table S1), which have been described as important regulators of oil body development (Kubo et al., 2018; Kanazawa et al., 2020; Romani et al., 2020; Romani et al., 2022). The patterns of expression were consistent with earlier published reporters (Romani et al., 2020; Takizawa et al., 2021) despite the shorter length of the promoters in our collection (30%, 49%, and 46% the length of the published promoters respectively). Having a comparable set of reporters allowed us to spot some differences between the expression patterns of each of them: *_pro_*Mp*ERF13* seems to be more active in oil body cells closer to the apical cell while *_pro_*Mp*R2R3-MYB02* is more evenly expressed in all oil body cells. In contrast, *_pro_*Mp*C1HDZ* expression is not restricted to only oil body cells (Romani et al., 2020). In addition, we observed that the reporters for Mp*BHLH34,* Mp*WRKY10,* Mp*REM2,* Mp*BHLH10,* Mp*TRIHELIX8,* Mp*ASLBD11,* and Mp*C2H2-8* displayed degrees of cell-type specificity, but their functions in Marchantia are largely unknown (Suppl. Table S1). Among them, Mp*C1HDZ*, Mp*R2R3-MYB2*, Mp*ERF13*, and Mp*WRKY10* mRNA were also shown to be specifically expressed in oil body cells in scRNA-seq (Wang et al., 2023). We also identified a set of promoters specifically active in rhizoid precursor cells (Fig. 5b). Of these, *_pro_*Mp*BHLH33/*Mp*RSL3* has been described before (Sauret-Gueto et al., 2020) and is very strongly expressed in all rhizoid cells (Fig. 5b). Some were active in the rhizoid precursors near the apical region but not in those located in the centre of the gemma (e.g., *_pro_*Mp*AP2L2 and _pro_*Mp*ERF2*), suggesting there are two populations of rhizoid precursor cells (proximal and distal) in the gemma (Fig 5b, Suppl. Table S1). Lastly, we observed a series of other promoters displaying seemingly random expression patterns that do not match any of the cell-types or expression domains that we have described here (Suppl. Table S1).

### Marker expression reveal the dynamics of cell fates in the stem cell zone

The availability of this prolific collection of highly precise cellular markers allows new approaches to visualizing the dynamics cell fates *in planta*. We followed the expression profile of a set of promoters active in the notch to better understand patterns of cell differentiation. We found five TFs reporters with high specificity for the SCZ (*_pro_*Mp*BZIP15, _pro_*Mp*BZIP7, _pro_*Mp*C2H2-26, _pro_*Mp*C2H2-22, _pro_*Mp*ERF20/LAXR, _pro_*Mp*CDC5*) at the gemma stage. The SCZ is composed of a central apical cell and a pair of immediate derivatives called sub-apical cells (Kohchi et al., 2021). During the first days of gemmaling development, it is possible to observe two stacked apical cells (Miller and Alvarez, 1965; Miller, 1966; Bowman, 2016). We followed the expression pattern of these candidates after the germination of gemmae and only *_pro_*Mp*ERF20/LAXR* remained expressed in the apical cells (Fig. 6, Suppl. Fig S2). In contrast, *_pro_*Mp*BZIP15, _pro_*Mp*BZIP7, _pro_*Mp*C2H2-26, _pro_*Mp*CDC5 and _pro_*Mp*C2H2-22* are expressed in a subset of differentiated cells after gemmae germination (Fig. 6, Suppl. Fig. S2). The expression of these reporters in sub-apical cells in the gemma provides evidence of the initiation of cell differentiation processes immediately adjacent to the stem cell.

**Figure 6.**
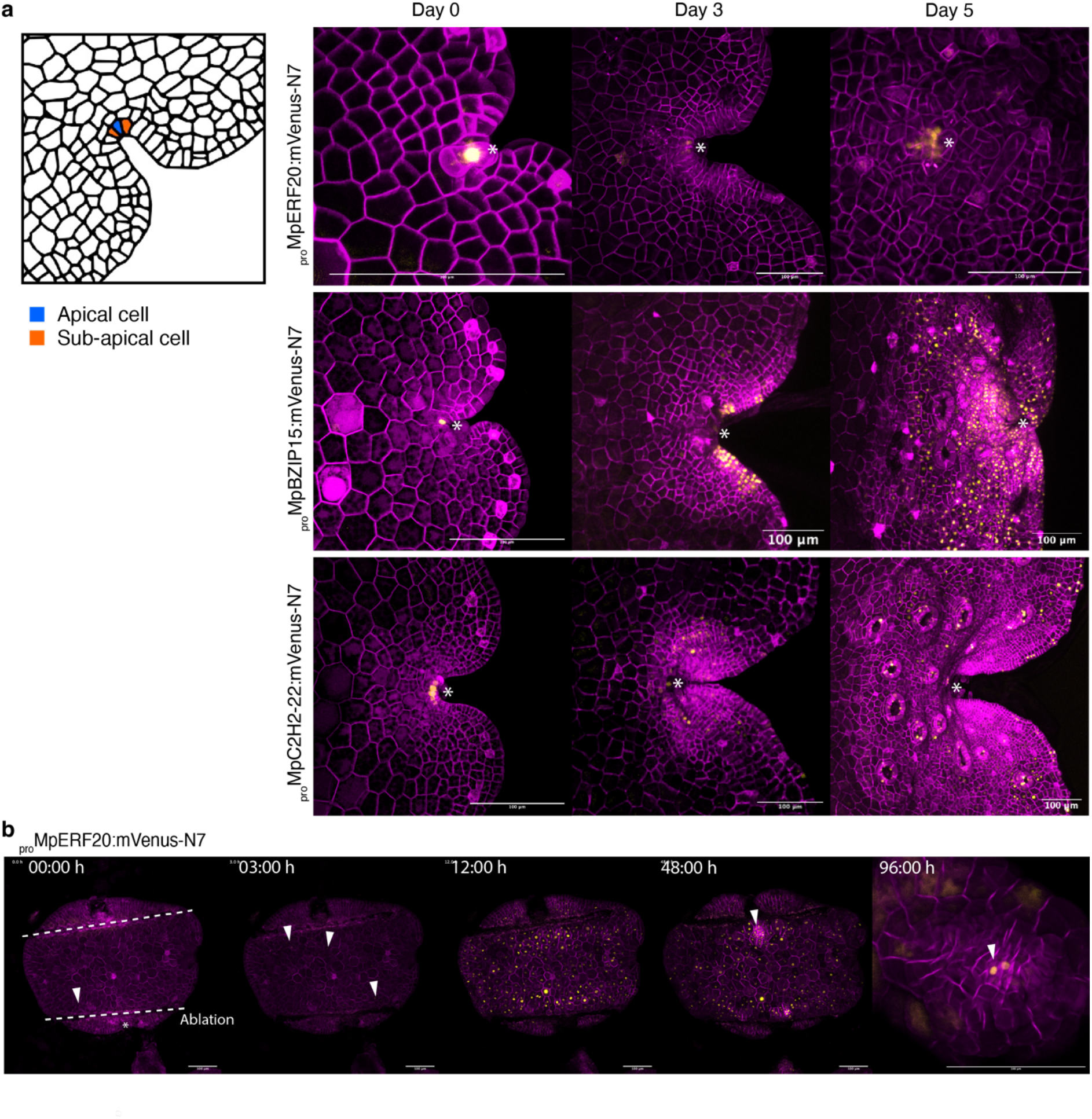
Dynamic expression of reporters in the SCZ. (a) A selection of promoters specifically active in the SCZ (*_pro_*Mp*ERF20/LAXR, _pro_*Mp*BZIP15, _pro_*Mp*C2H2-22)*. Cell types of the SCZ are shown on the left. Confocal images of the gene of interest (yellow) and a constitutive plasma membrane marker (magenta, *_pro_*Mp*UBE2:mScarlet-Lti6b*). Asterisks point the apical notch. (b) Time lapse of *_pro_*Mp*ERF20/LAXR* expression after laser ablation of the notches and until re-establishment of the new SCZ. Ablated regions are marked as dotted lines. Arrows point the first cells with signal and the forming apical notch (see also Supplemental Movie S1). Scale bars = 100 μm. Gene IDs: Mp*ERF20/LAXR =* Mp5g06970, Mp*BZIP15 =* Mp1g03580, Mp*C2H2-22 =* Mp4g11030.

We verified the expression of Mp*ERF20/LAXR* transcript by *in situ* hybridisation. (Suppl. Fig. S4). Tissue-specific expression of Mp*ERF20/LAXR* in the SCZ was confirmed, however mRNA transcript signal corresponds to a larger area than it was observed in the transcriptional reporter. It was recently shown that Mp*ERF20/LAXR* plays a fundamental role in regeneration, has the capacity to induce cellular reprogramming to generate undifferentiated cells and it is a sufficient to generate new apical stem cells (Ishida et al., 2022). After the ablation of the notches in the gemma, a strong response of *_pro_*Mp*ERF20/LAXR* is induced in the whole tissue after just 5 hours consistently with data from RNA-seq experiments after ablation (Ishida et al., 2022). A previous longer version (4.3 kb vs 1.8 kb) of this promoter also displayed similar induction after ablation (Ishida et al., 2022). Following the induction of *_pro_*Mp*ERF20/LAXR*, cells start dividing and de-differentiate until a new apical region is formed. Subsequently, the expression activity of *_pro_*Mp*ERF20/LAXR* diminishes in epidermal cells and only remains in the new SCZ (Figure 6, Supplemental movie S1).

### Dynamic expression of reporters during gemmaling development

Outside of the SCZ, we identified 20 TF promoter-driven reporters specifically expressed in the DDCZ. Interestingly, this later group of TFs also include stress related genes such as Mp*ABI3a*, Mp*MYCY*, Mp*HY5* (Clayton et al., 2018; Eklund et al., 2018; Penuelas et al., 2019), suggesting that stress signal transduction pathways are specifically active in the DDCZ at this stage of gemma development. We found only 9 TF reporters specific to the TZ, and all of them are also active in the DDCZ and SCZ. This logic is followed by other TFs expressed in the PZ. Altogether, the Marchantia meristem is characterised by more than 200 TFs active in the SCZ and this number diminishes as cells mature and are displaced distally from the apical growth direction.

Two types of expression pattern do not follow a regular profile along the apical axis and are not associated with known specialised cells. The first corresponds to cells around the perimeter of gemmae that we called “border cells”. Such cells were not well described in the literature. In a transverse section, the border cells form a layer of 2-3 cells at the margins of the gemma. Among the promoters observed, Mp*NAC2* and Mp*ARF2* show higher specificity for expression in border cells. A similar expression pattern was shown before by using a knock-in reporter of Mp*ARF2* (Kato et al., 2020b). We observed the expression of both genes after gemma germination (Fig. 7a, Suppl. Fig. S3a) and the expression maximum migrates from the border to the CZ after 2 days. We believe the border expression pattern could be associated with the establishment of abaxial/adaxial polarity or auxin accumulation during gemma formation. This interpretation is supported by the role of Mp*ARF2* and auxin signalling in gemmae development (Rousseau, 1953; Eklund et al., 2015).

**Figure 7.**
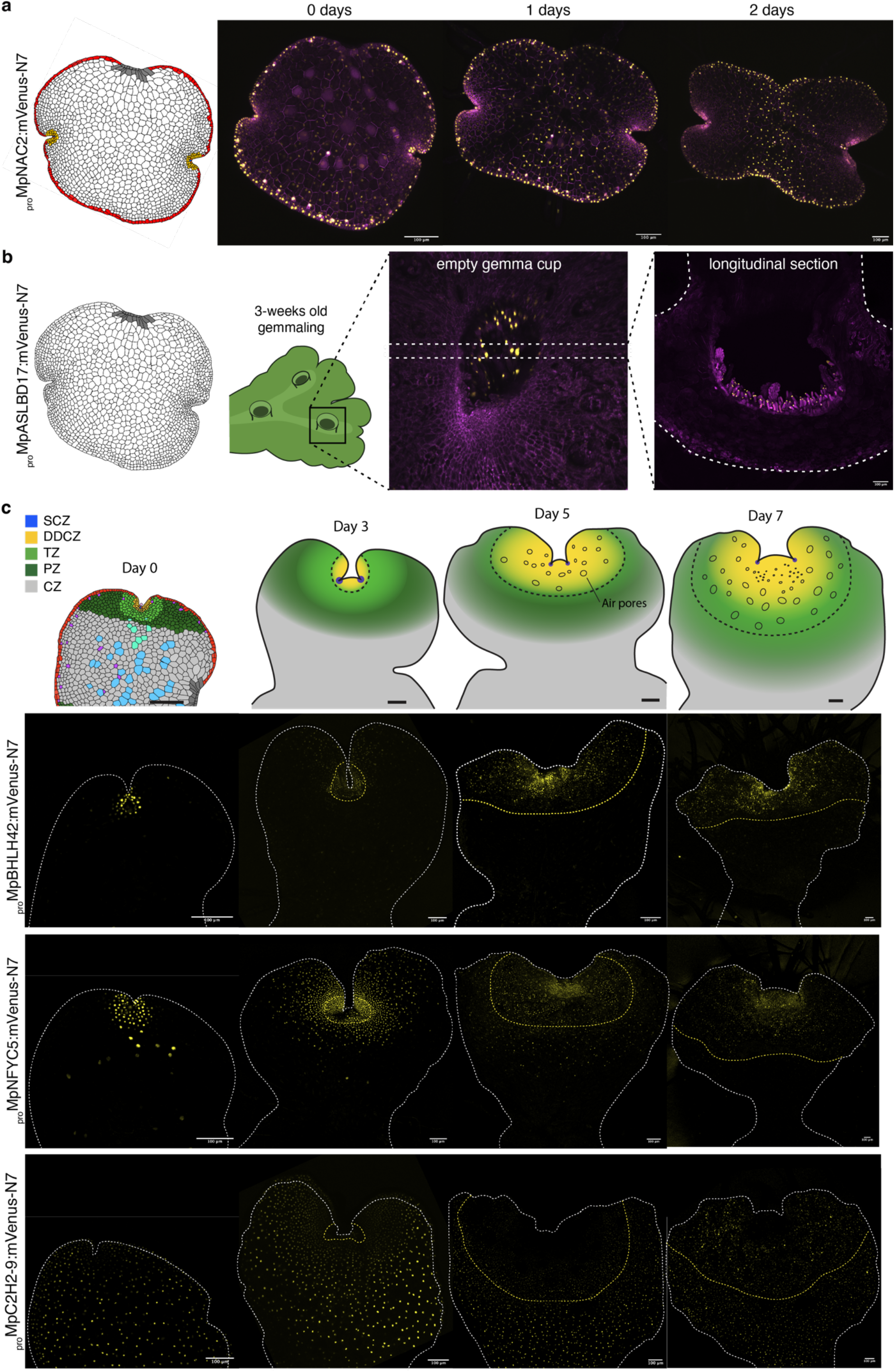
Promoter activity during gemmaling development. (a) Schematic representation of the gemma border and time-course of expression for a representative cell-type specific marker (*_pro_*Mp*NAC2*). (b) Schematic representation of the attachment point of the gemma and expression of a representative cell-type specific marker (*_pro_*Mp*ASLBD17*) in a gemma cup in a mature thallus (view from the top and cross-section). Confocal images of the gene of interest (yellow) and a constitutive plasma membrane marker (magenta, *_pro_*Mp*UBE2:mScarlet-Lti6b*). (c) Schematic models of expression domain dynamics during the first days of gemmaling development, with (below) examples of confocal images of time-courses of fluorescent reporters (*_pro_*Mp*BHLH42, _pro_*Mp*NFYC5, _pro_*Mp*C2H2-9*) illustrating the different expression domains. The dashed line represents the boundary between the mature epidermis and the supportive tissue of the gemmae. Scale bar 100 μm. Gene IDs: Mp*NAC2 = Mp6g02590,* Mp*ASLBD17 = Mp8g09250,* Mp*BHLH42 = Mp5g09710,* Mp*NFYC5 = Mp1g16880,* Mp*C2H2-9 = Mp7g09260*.

The second special expression domain corresponds to a group of elongated cells referred to in the literature as the “attachment point” (Solly et al., 2017) which correspond to the cells connected to the cup base before the detachment of the gemma from the stalk cell. These (Kato et al., 2020a). _pro_Mp*ASLBD17* is the best reporter with high specificity for the attachment cells (Fig. 5). After gemma germination, _pro_Mp*ASLBD17* signal remain in the attachment but signal diminishes (Suppl. Fig. S3). Interestingly, Mp*ASLBD17* is also active in the base of the cup in both gemma initials and mucilage cells (Fig. 7b). These terminal cells do not divide after germination suggesting this cell identity could be a remnant of interaction between the gemma and cup. Later in development, *_pro_*Mp*ASLBD17* is also strongly expressed in slime papillae (Suppl. Fig. S3b). This is consistent with the notion that the mucilage papillae, slime papillae, and gemma initials are all homologous cell types with similar genetic programs (Proust et al., 2016).

We followed the expression patterns of 27 other promoters active in the different expression domains in the notch across the course of vegetative development in Marchantia gemmalings for 7 days. During this developmental period, gemmaling start maturating and proliferating and undergo drastic morphological changes. Still, most patterns remained consistent (19/27) with the pattern of cell divisions (Fig. 7c). Examples of expression patterns in 0, 3, 5 and 7 days-old are shown in Figure 7c. After the first 2 days of growth, cells rapidly expand and form a mature epidermis while the first bifurcation of the thallus takes place. Proximal rhizoids (Fig. 5a) of the dorsal surface can still undergo cell divisions and de-differentiate into epidermal cells, while distal rhizoids are committed to elongate even at the dorsal surface. The mature thallus is characterised by the complete formation of air chambers and air pore structures (Shimamura, 2016). These structures are formed by a very precise pattern of cell divisions that occur very close to the SCZ and form a boundary between the mature thallus and the gemmae epidermis visible after 3-4 days. The DDCZ drastically expands during the first days and covers most of the newly formed mature thallus, displacing the TZ and PZ (Fig. 7C). This contrasts with TF promoters expressed in the SCZ of the gemma which remain limited to sub-domains of the mature thallus (Fig. 6). The DDCZ, TZ, and PZ maintain a high rates of cell expansion and division during the first days (Boehm et al., 2017; Ishida et al., 2022) but only the DDCZ is active during the differentiation of cells. Both the TZ and PZ expand to form the boundary and heart-shaped morphology that separate both apical notches, acting as a supportive tissue to the forming mature epidermis (Fig. 7c). The CZ remains unaltered while the rhizoid precursors in the dorsal region de-differentiate. It is only after 5-7 days that the DDCZ forms a gradient of expression focused on the SCZ forming a boundary between developing and mature air pores (Fig. 7c). This structure is repeated in a similar pattern during vegetative growth (Solly et al., 2017). We synthesized these observations and expanded our model of expression domains to later developmental stages (Fig. 7c).

The mature thallus is later characterised by the presence of air pores and gemma cups. We also observed that other promoters showing tissue specific expression in organs in the mature thallus, such as gemma cup (*_pro_*Mp*NAC1 and _pro_*Mp*ERF11*) and air pores (*_pro_*Mp*C3H8, _pro_*Mp*CCAAT-NFYC4*), are also active in the DDCZ of the developing gemma (Fig. 8). Among them, Mp*CCAAT-NFYC4* was also found to be air pore specific in scRNA-seq experiments (Wang et al., 2023). These observations are in agreement with classical morphological models of cell differentiation in bryophytes where most cell differentiation processes occur in the cells surrounding the apical region (APOSTOLAKOS et al., 1982; Bowman, 2016; Shimamura, 2016).

**Figure 8.**
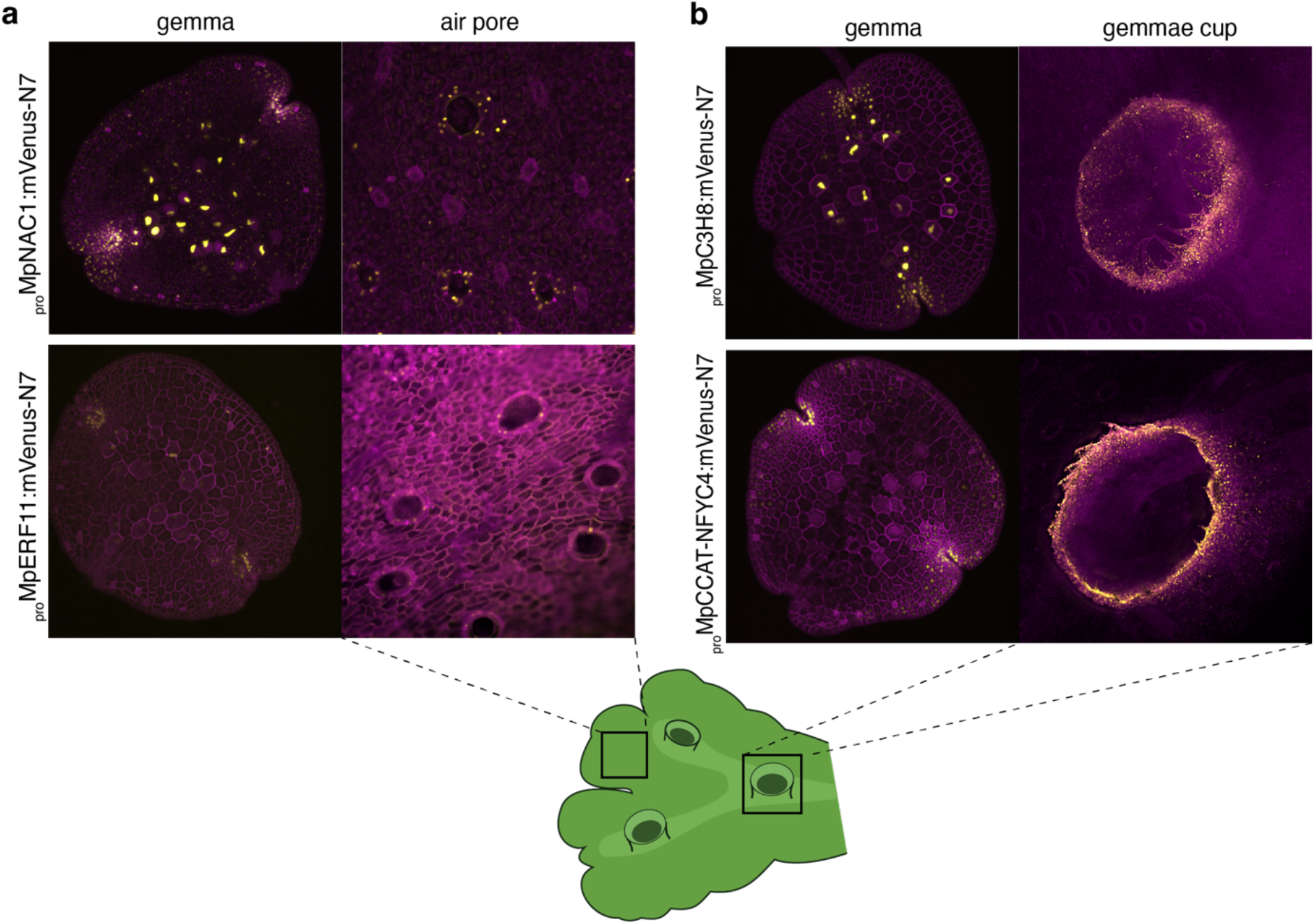
Promoters specifically active in mature thallus tissues. (a) Expression pattern of reporters with specific expression in air pores (*_pro_*Mp*C3H8, _pro_*Mp*CCAAT-NFYC4*) and (b) gemma cups *(_pro_*Mp*NAC1 and _pro_*Mp*ERF11)*. Schematic representation of a Marchantia adult plant and the correspondence to images shown. Gene IDs: Mp*C3H8 = Mp2g05060,* Mp*CCAAT-NFYC4 = Mp1g01960,* Mp*NAC1 = Mp2g07720,* Mp*ERF11 = Mp7g17020*.

## DISCUSSION

We built and tested a comprehensive library of promoters derived from the genes of regulatory TFs in Marchantia. The promoter parts are of relatively compact size with standardised modular format to allow simple DNA engineering. These reporters can be easily reused and combined to recognise virtually any cell-type in Marchantia, providing a toolset that rivals any other plant system.

We have used these promoters to systematically map patterns of gene expression during early gemmaling development in Marchantia. We exploited nuclear-localised fluorescent cell markers and the regular cellular architecture of gemmae to normalise and compare patterns of gene expression with cellular resolution. These could be registered using microscopic features of cellular anatomy and compared with published knowledge of cellular differentiation to enable construction of a stereotypical map of cell states in the Marchantia gemma. This atlas will provide a guide for further use of the promoter collection, and a template for more detailed studies of the interactions between genome and cellular development in Marchantia.

The promoter activities have some limitations in accurately reflecting the transcriptional patterns of the corresponding endogenous genes. For example, they may be missing important downstream or upstream regulatory regions, alterations due to domestication, and post-transcriptional regulatory mechanisms associated with the native transcripts. The former, was shown to be important for several developmental regulators in Marchantia (e.g., Mp*RSL1*, Mp*FGMYB* (Honkanen et al., 2018; Hisanaga et al., 2019)). Nevertheless, we found broad and consistent correlations between the observed patterns of promoter activity and independently measured levels and distribution of transcripts, and documented properties of longer versions of the promoters from the literature. Our approach is complementary to other transcriptomic efforts to understand Marchantia development. Moreover, it could capture precise features of cellular organisation and gene regulation in the apical meristem that were not discernible by time-resolved scRNA-seq (Wang et al., 2023). Further, these promoters can drive expression of fluorescent proteins to deliver spatially precise and sensitive markers for visualising the dynamics of cell states in living tissues.

Reconstructing the evolution of morphological traits requires defining the relationship between tissues and cell types and how genetic programs evolved (Delaux et al., 2019; Zeng, 2022). Previous models suggested that the vegetative gametophyte meristem of bryophytes is analogous or homologous to the vegetative sporophyte meristem in tracheophytes (vascular plants), both as a deeply conserved trait or by the co-option of several TFs from one generation to the other (Bowman et al., 2019). To reconstruct the history of the evolution of meristems in land plants, the expression patterns of TFs play a crucial role. Looking only for conserved factors across embryophytes may have generated constraints in the comparisons between the functional architectures of these two forms of multicellular polar growth. Our approach of testing a near-complete collection of TF reporters has the potential to revisit this question, sidestepping selection bias.

Morphological studies suggest that stem cells of the vegetative body of bryophytes are comprised of single apical cell (Menand et al., 2007; Shimamura, 2016; Suzuki et al., 2020). This simple structure is likely the ancestral state of the land plant meristem, while the more complex meristem observed in vascular plants is likely a derived trait (Harrison and Morris, 2018; Fouracre and Harrison, 2022). Our observations provide genetic evidence for the identity of such cells in Marchantia. Mp*ERF20/LAXR* is expressed in the centre of the SCZ (as verified by the fluorescent reporter and *in situ* hybridisation) and accompanied by sub-apical cells where other TFs are specifically expressed (Mp*BZIP15,* Mp*BZIP7,* Mp*C2H2-26,* Mp*C2H2-22, MpCDC5*). In addition, we found a set of TF promoters active in the DDCZ that completes the arrangement of cells forming the Marchantia notch, that constitute the building blocks of Marchantia vegetative development.

Thus, there appears to be a hierarchical order to the patterns for gene expression in the Marchantia thallus. Many TFs are expressed in the SCZ and expression patterns are progressively pruned along the longitudinal axis as distal daughter lineages take up specific cell fates (Fig. 4d). However, we also observed complex gene expression patterns which are active in broad domains but excluded from specific cell-types (e.g., *_pro_*Mp*ERF21, _pro_*Mp*BZR2, _pro_*Mp*BBX3*) that could also be important for developmental processes.

The classical model of stem cell organization in the sporophyte of vascular plants involve *WUSCHEL (WUS/WOX), Class I KNOX (KNOX1), Class III HD-ZIP (C3HDZ), INTEGUMENTA/PLETHORA/BABYBOOM (APB), SCARECROW (SCR), SHORTROOT (SHR)* and *HAIRY MERISTEM (HAM) TFs*. In Marchantia, the reporters for Mp*WOX*, Mp*APB*, Mp*KNOX1*, and Mp*C3HDZ* are not specific to an analogous region of the apical notch in the Marchantia gametophyte (Suppl. Table S1). This is in line with functional evo-devo studies in bryophytes showing that Mp*WOX* does not play a critical role in the gametophyte of Marchantia (Hirakawa et al., 2020), that Mp*KNOX1* only participates in the sporophyte generation (Sano et al., 2005; Sakakibara et al., 2008; Dierschke et al., 2021; Hisanaga et al., 2021), and C3HDZ mutant does not affect the gametophytic meristem in the model moss *P. patens* (Yip et al., 2016). As observed in other cases, the function of TFs could be only conserved in the sporophyte generation (Romani and Moreno, 2021). In the case of GRAS TFs such as HAM and SCR, they seem to play a prominent role in the gametophytic stem cell organization in Physcomitrium, but orthologues for some do not exist in Marchantia (Beheshti et al., 2021; Ge et al., 2022; Ishikawa et al., 2023).

The singular set of TFs expressed in the SCZ is largely unrelated to known TFs associated with meristem organization in other species. For example, Mp*BZIP15* has no true orthologue in angiosperms (Bowman et al., 2017) and characterized C2H2 TFs are largely associated with stress responses (Han et al., 2020). Interestingly, *CDC5* has been associated with shoot apical meristem organization in Arabidopsis upstream of *STM* and *WOX* and loss-of-function plants are embryo lethal, but its expression is not meristem specific (Lin et al., 2007). As a possible exception, in *Arabidopsis*, At*ESR1/DRN* the orthologue of Mp*ERF20,* was described to be involved in regulation of the shoot apical meristem organization and regeneration, suggesting that this role could be conserved across land plants, or co-opted in the opposite generation (Banno et al., 2001; Kirch et al., 2003; Ikeda et al., 2021). However, unlike Mp*ERF20*, At*ESR1/DRN* is expressed in the leaf primordia and not in the stem-cell zone (Kirch et al., 2003).

In summary, the evidence presented here supports the notion that GRNs governing the formation of an apical meristem in the vegetative body of bryophytes and embryophytes are not analogous. One scenario is that both forms of multicellular polar growth evolved to a large degree independently in contrasting generations. The fact that the only conserved factor is associated with regeneration, indicates that the bryophyte meristem GRNs may be built on top of an ancestral capacity of ancestral land plants to regenerate. On the other hand, the more complex body plans of vascular plant may have recruited *de novo* GRNs during evolution to support organ development and more sophisticated patterning. In contrast, most of the differentiation events in Marchantia development are observed immediately after formation of the first derivatives of the apical cell (DDCZ) and there is not a comparable peripheral zone as in the sporophyte of vascular plants. Nevertheless, other aspects of the molecular machinery regulating the meristem formation and maintenance, such as: peptide signalling, such as auxin biosynthesis and polar transport, and cytokinin signalling; seem to work in a similar fashion in both forms of vegetative body (Whitewoods et al., 2018; Aki et al., 2019; Hirakawa et al., 2019; Blazquez et al., 2020; Hirakawa et al., 2020; Kato et al., 2020b; Bowman et al., 2021). Future work on hormone control of growth and their interaction with TFs in bryophytes and streptophyte algae will help to fill the gaps in how the cell types are defined and maintained across development.

This atlas of TF expression patterns will provide a valuable resource for the plant science community. As we showed for the case of the air pores and cups, there is a strong potential to find tissue-specific promoters to Marchantia tissues in other developmental stages not covered here. We expect this collection of promoter will help to accelerate studies in Marchantia for a wide range of applications: markers for cell identities, ratiometric quantification (Federici et al., 2012), isolation of nuclei tagged in specific cell types (INTACT)(Deal and Henikoff, 2011), cell-type specific expression, among many other functional genomics and synthetic biology applications.

## ACKNOWLEDGMENTS

We thank the Marchantia evo-devo community for useful discussion. We thank Nicola Patron and the Earlham institute Biofoundry for assistance with the automated cloning. This work was funded as part of the BBSRC/EPSRC OpenPlant Synthetic Biology Research Centre Grant BB/L014130/1 to J.H., BBSRC BB/F011458/1 for confocal microscopy, BBSRC BB/T007117/1 to J.H, and Australian Research Council grants DP210101423, CE200100015 to J.L.B.

## METHODS

### Plant Material and Growth Conditions

*Marchantia polymorpha subs. rudelaris* accessions *Cam-1* (male) and *Cam-2* (female) were used in this study (Delmans et al., 2017). Under normal conditions, plants were grown on solid 0.5× Gamborg B-5 basal medium (Phytotech #G398) at pH 5.8 with 1.2% (w/v) agar micropropagation grade (Phytotech #A296), under continuous light at 21 °C with light intensity of 150 μmol/m^2^/s. For spore production, plants were grown in Microbox micropropagation containers (SacO_2_) in long day conditions (16 h light/8 h dark) under light supplemented with far-red light as described (Sauret-Gueto et al., 2020).

### Synthesis of L0 parts

5’UTR and promoter regions from genes were extracted from Marchantia genome version 5.1 (Montgomery et al., 2020) primary transcripts. DNA sequences were domesticated to remove internal BsaI and SapI sites. The sequences of synthetic L0 parts used in this work is available in Supplemental Table S1. L0 parts were synthesised either by GENEWIZ or Twist Bioscience following the standard syntax for plant synthetic biology with PROM5 or PROM and 5UTR overhangs and cloned into the plasmid pUAP1 (Addgene #63674) (Patron et al., 2015) by recombination. Promoter sequences with repeated Ns in first 1000bp or 5UTR longer than 3 kbp, were omitted. Additional L1 and L0 parts were obtained from the OpenPlant toolkit (Sauret-Gueto et al., 2020)(Supplemental Table S1).

### Plasmid assembly

L1 and L2 plasmids were constructed using Loop Assembly as described before (Pollak et al., 2019) with the L0 and L1 parts described in Supplemental Table S1. For one-step assembly of L3 plasmids, a new acceptor (pBy_01) was built using NEBuilder HiFi DNA Assembly Master Mix (New England Biolabs, NEB #E2621). Four fragments were amplified by PCR using the Q5 High-Fidelity DNA Polymerase (NEB #M0492) and purified using Monarch PCR & DNA Cleanup kit (NEB #T1030). The *proUBE2:mTurquoise-N7; proUBE2:mScarlet-Lti6b; pro*Mp*WRKY10:mVenus-N7* plasmid was used as a template, with primers Fw1 (5’-acataacgaattgctcttcaagattagccttttcaatttcagaaagaatg-‘3) and Rv1 (5’-ggtctctctccctccctccttgctagcgatc-‘3), Fw2 (5’-cctgtcgtgcggtctcaaatggtgagcaagggcgaggagc-‘3), Rv2 (atctcgaatccgacggccacgcggcatg-‘3), Fw3 (5’-gtggccgtcggattcgagatccaccgag-‘3), Rv3 (5’-cctgtcagaattgctcttcaatctggattttagtactggattttg-‘3); and pCsA as template with primers Fw4 (5’-aaggagggagggagagagaccagcttgtctgtaagcggatg-‘3) and Rv4 (5’-catttgagaccgcacgacaggtttcccgac-‘3). The full-length of the final construct was verified by sequencing using the Oxford Nanopore technology (SNPsaurus LLC). The acceptor pBy_01 was used to assemble using BsaI and L0 corresponding to PROM5 or PROM and 5UTR parts as in Supplemental Table S1. Type-IIS cloning was performed as described previously (Cai et al., 2020) using a Master Mix containing 10% (v/v) 10× T4 DNA ligase buffer (NEB #M0202), 2.5% (v/v) 1 mg/mL bovine serum albumin (NEB #B9200S), 5% (v/v) T4 DNA ligase at 400 U/μL (NEB #M0202), 5% (v/v) BsaI at 20 U/μL (NEB #R3733), 10% (v/v) acceptor at 40 ng/uL, 20% (v/v) pre-mixed L0 parts (∼100 ng/μl), and water to a final volume of either 2 μL for the acoustic liquid handling robot (Labcyte Echo 550, Beckman) or 5 μL for manual handling. Cycling conditions were 26 cycles of 37 °C for 3 min and 16 °C for 4 min. Termination and enzyme denaturation: 50 °C for 5 min, and 80 °C for 10 min. 15 μL of TOP10 chemically competent E. coli cells were transformed using the assembly reaction and plated on LB-agar plates containing 50 μg/mL kanamycin and 40 μg/mL of 5-bromo-4-chloro-3-indolyl β-D-galactopyranoside (X-Gal). The presence of the correct insert was confirmed by restriction XhoI digestion (Thermo Scientific #FD0694) and Sanger sequencing using primers Fw5 (5’-tactcgccgatagtggaaacc) and Rv5 (5’-aagcactgcaggccgtagcc-‘3).

### Agrobacterium mediated transformation

Marchantia spores were sterilised as previously described (Sauret-Gueto et al., 2020). A modification of the published Agrobacterium-mediated protocol for transformation in multi-well dishes was used (Ishizaki et al., 2008; Sauret-Gueto et al., 2020). Briefly, *A. tumefaciens* (GV3103) were transformed using a miniaturised freeze-thaw method (Weigel and Glazebrook, 2006) and plated in six-well plates with LB-agar plus kanamycin (50 mg/ml), rifampicin (50 mg/ml), and gentamycin (25 mg/ml) and grown for 3 days at 29°C. Spores were grown on solid 0.5× Gamborg B-5 media for 5 days and dispensed in 6-well plates containing 4 mL of liquid 0.5× Gamborg B-5 plus supplements: 0.1% N-Z amino A (Sigma #C7290) 0.03% (w/v) L-glutamine (Alpha Caesar #A14201) 2% (w/v) sucrose (Fisher Scientific #10634932), and 100 μM acetosyringone. A single colony of *Agrobacterium* transformed with the plasmid of interest was scooped and inoculated the spore culture. The 6-well plate was then placed on a shaker at 120 rpm for 2 days at 21 °C with continuous lighting (150 μmol/m^2^/s). For each well, the sporelings were washed with 25 mL of sterile water and plated on solid 0.5× Gamborg B-5 media supplemented with 0.5% (w/v) sucrose plus 100 μg/mL cefotaxime (Apollo Scientific, #BIC0111) and hygromycin 20 μg/mL (Invitrogen, #10687010). Plants were grown in normal conditions for 10 days and transferred to a new selection plate for another 12-14 days until cups with gemmae are formed.

### Laser Scanning Confocal Microscopy

Images of Marchantia were acquired on a Leica SP5 confocal microscope upright system equipped with Argon ion gas laser with emitted wavelengths of 458, 476, 488 and 514 nm, 405 nm diode laser, 594 nm HeNe laser, 633 nm HeNe laser, and 561 DPSS laser. For higher resolution and time lapse studies, images were acquired on a Leica SP8X spectral confocal microscope upright system equipped with a 460−670 nm super continuum white light laser, 2 CW laser lines 405 nm, and 442 nm, and 5 Channel Spectral Scanhead (4 hybrid detectors and 1 PMT). For slides, imaging was conducted using either a 10× air objective (HC PL APO 10×/0.40 CS2), a 20× air objective (HC PL APO 20×/0.75 CS2). When observing fluorescent protein with overlapping emission spectra, sequential scanning mode was selected. Excitation laser wavelength and captured emitted fluorescence wavelength window were as follows: for mTurquoise2 (442 nm, 460−485 nm), for eGFP (488 nm, 498−516 nm), for mVenus (514 nm, 527−552 nm), for mScarlet (561 nm, 595−620 nm), and for chlorophyll autofluorescence (633, 687−739 nm).

When imaging time-courses, plants grown under normal culture conditions in small petri dishes, removed the lid for imaging, and returned the plants to the growth chamber and imaged as described above. For live imaging, six stacked Gene Frames (ThermoFisher #AB0578) were placed on a glass slide and filled halfway with molten Gamborg B-5 agar medium. Plants were then places on the solidified agar surface and meristems were removed using a Laser Microdissection Leica LMD7000. Samples were mounted in perfluorodecalin (Littlejohn et al., 2010) with a glass coverslip on top. The slides were then continuously imaged on the Leica SP8X confocal microscope for 1−4 days.

### Analysis of Public RNA-Sequencing Data

Transcripts per million (TPM) values were extracted from Marpolbase Expression database (Kawamura et al., 2022). Sample accessions DRR284685 and DRR284686 (Mizuno et al., 2021) were used to compare reporter expression patterns with RNA-seq. Data was subsequently analysed with R. Hyperbolic arc-sin was calculated for each corresponding transcript (*base* package v4.1.3) and plotted with the density function (*stats* package v4.1.3).

### Image analysis and clustering

Image processing was performed in Fiji (Schindelin et al., 2012) to perform maximum intensity projections of the Z-stacks. For fluorescence intensity analysis, background was subtracted with parameters by default, images were rotated to align the notches in the X-axis, and the histogram was done using the plot profile function of the mVenus channel covering the entire gemma, using the chlorophyll channel as a reference. Raw intensity data and distance of the notches was exported for further analysis in R. The *smooth.spline* function (spar=0.4) was used to reduce noise from cell-to-cell signal, and *approxfun* function from the *stats* package was used to interpolate the distance from the start to the first notch, and then to the second and end of the plot using fixed values. The average distance values of all images taken was used as a reference to align all profiles. Intensity was normalised to the maximum value. The *hclust* and *cutree* functions from *stats* package were used to perform the clustering and extract the groups. The *pheatmap* function for *ComplexHeatmap* package v2.10.0 (Gu et al., 2016) was used to plot the heatmap. For calculating the mode, the *mlv* function from *modeest* package v2.4.0 with the Grenander method (Grenander, 1965). Default parameters were used, and plots were made using the *base* package v4.1.3 unless specifically stated.

### In situ hybridisation

Mp*ERF20* coding sequence was amplified from cDNA using primers MpERF20 cds in situ F (5’-GTACAAAAAAGCAGGCTCCGCGGCCGCatggtggggagg-3’) and MpERF20 cds in situ R (5’ GTACAAGAAAGCTGGGTCGGCGCGCCttacatgagtgggggaactaaaagaagagt-3’) and seamlessly cloned using NEBuilder HiFi DNA Assembly (New England Biolabs, #E5520) into pENTR-D linearized with *Not*I/*Asc*I. *M. polymorpha* ssp *ruderalis*, ecotype MEL, tissue fixation, embedding, sectioning, and hybridization with digoxigenin (DIG)-labeled antisense RNA probes were performed according to (Zachgo, 2002). Microscopic slides were observed using an Axioskop 2 mot plus (Zeiss) microscope and photographed using AxioCam HRc and AxioVision software.

---

**Supplemental Figure S1.**
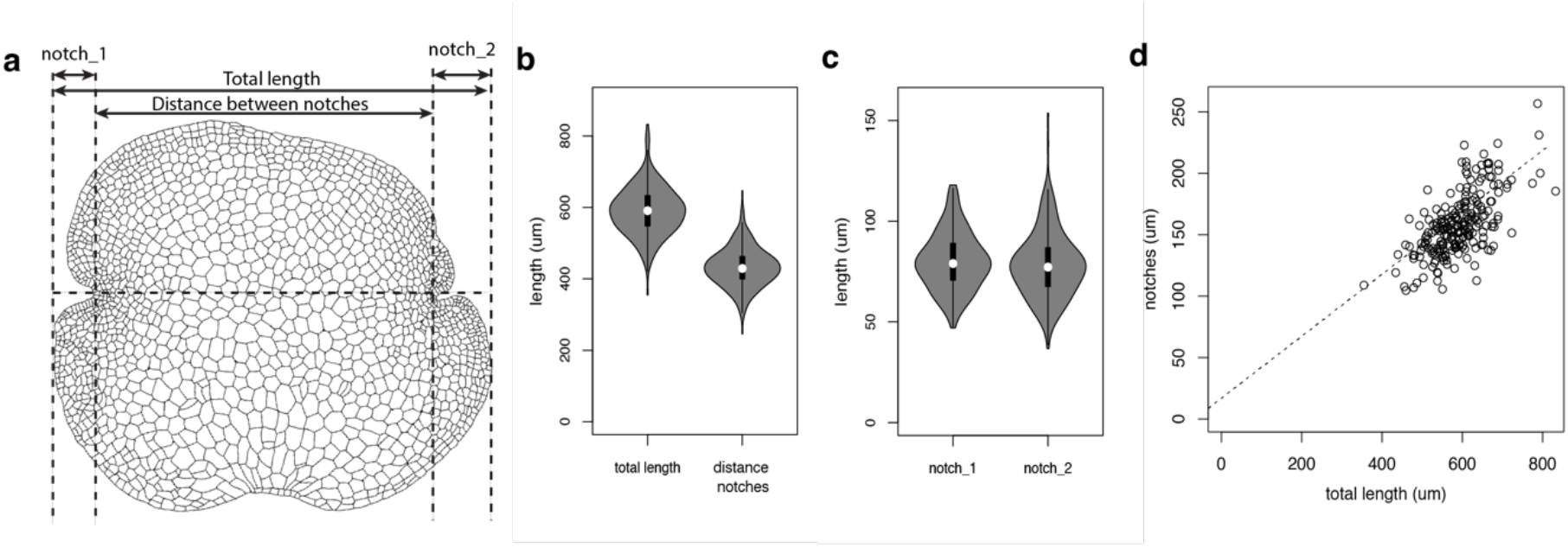
Variability of the Marchantia gemma dimensions. (a) Schematic drawing of Marchantia gemma dimensions. (b) Distribution of total length and distance between notches. (c) Distribution of length between notches and gemma border. (d) Correlation between total length and distance from notch to border.

**Supplemental Figure S2.**
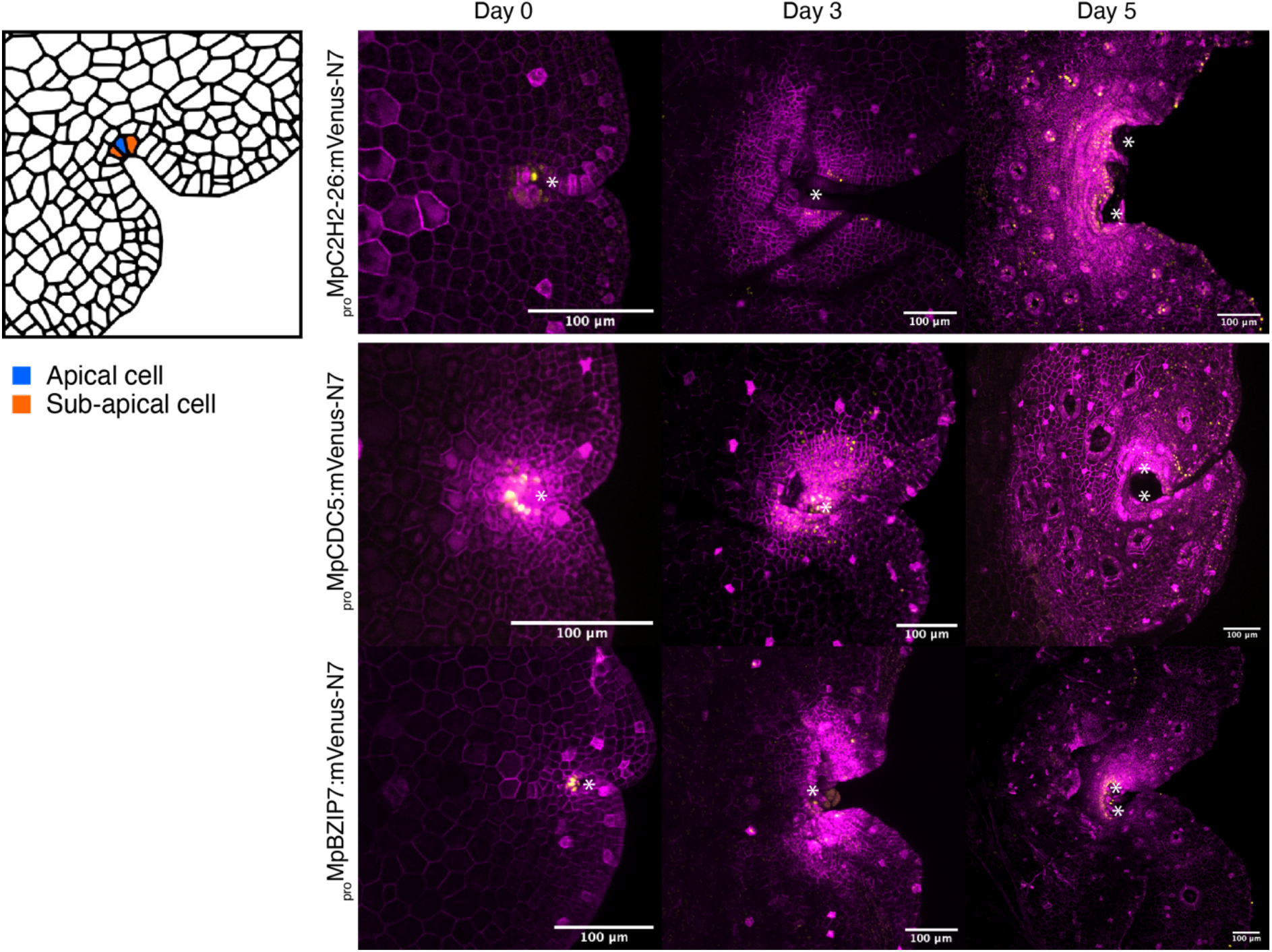
Dynamic activity of additional TF promoters reporters specific to the SCZ (*_pro_*Mp*C2H2-26, _pro_*Mp*CDC5, _pro_*Mp*BZIP7) driving expression of mVenus-N7 nuclear-localised fluorescent protein).* Cell types of the SCZ are shown on the left. Confocal images of the gene of interest (yellow) and a constitutive plasma membrane marker (magenta, *_pro_*Mp*UBE2:mScarlet-Lti6b*). Asterisks mark the apical notch. Scale bars = 100 μm. Gene IDs: Mp*C2H2-26 = Mp8g14220,* Mp*CDC5 = Mp1g10310,* Mp*BZIP7 = Mp3g04360*.

**Supplemental Figure S3.**
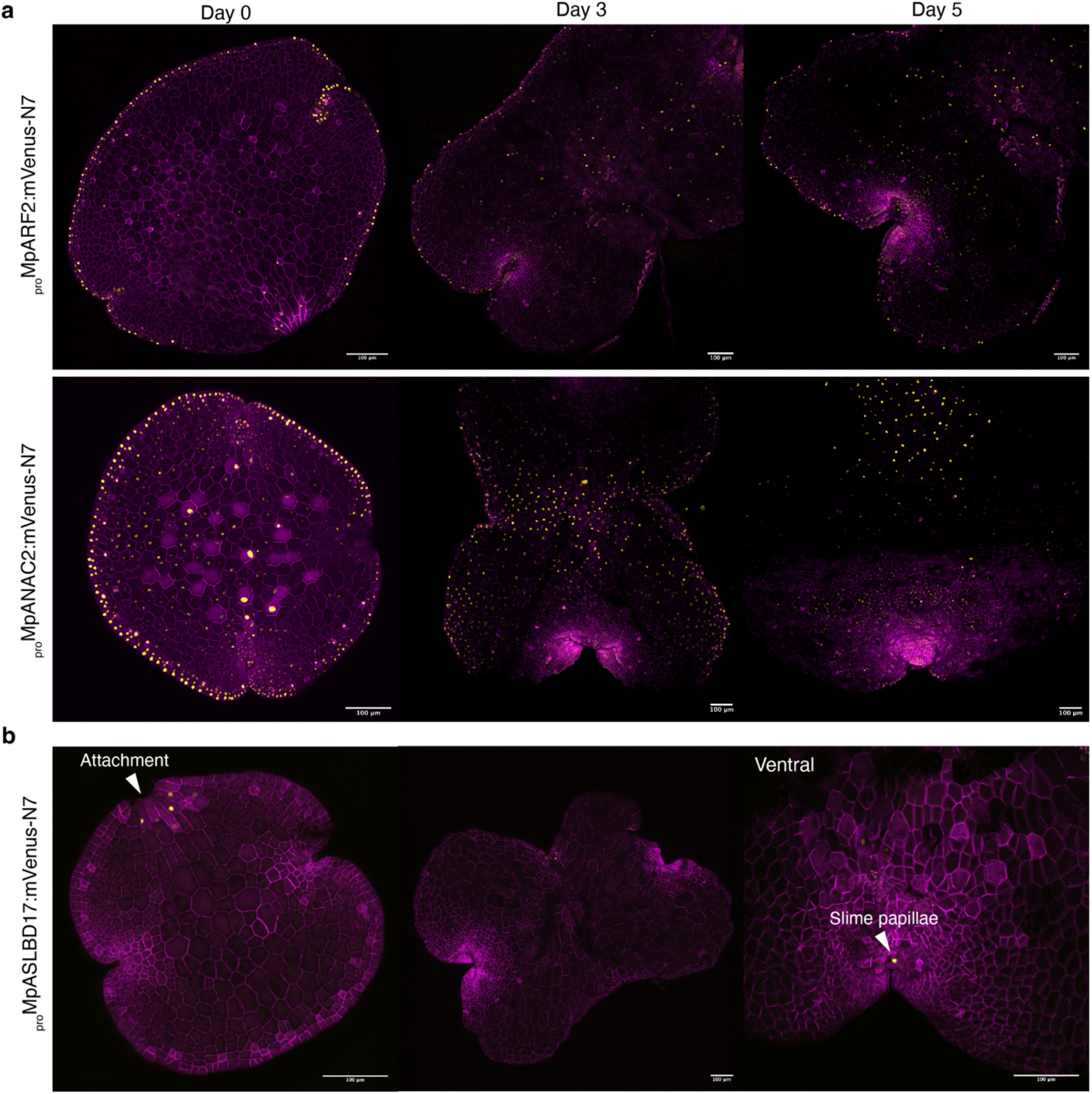
(a) Time-course images of gemmaling development in lines with expression of fluorescent reporters specific to the border (*_pro_*Mp*ARF2 _pro_*Mp*NAC2)*. (b) Time-course images of gemmaling development in lines with expression of fluorescent reporters specific to the attachment (*_pro_*Mp*ASLBD17*). The attachment and slime papillae are highlighted. Confocal images of the gene of interest (yellow) and a constitutive plasma membrane marker (magenta, *_pro_*Mp*UBE2:mScarlet-Lti6b*). Scale bar 100 μm. Mp*ARF2 = Mp4g11820,* Mp*NAC2 = Mp6g02590,* Mp*ASLBD17 = Mp8g09250*.

**Supplemental Figure S4.**
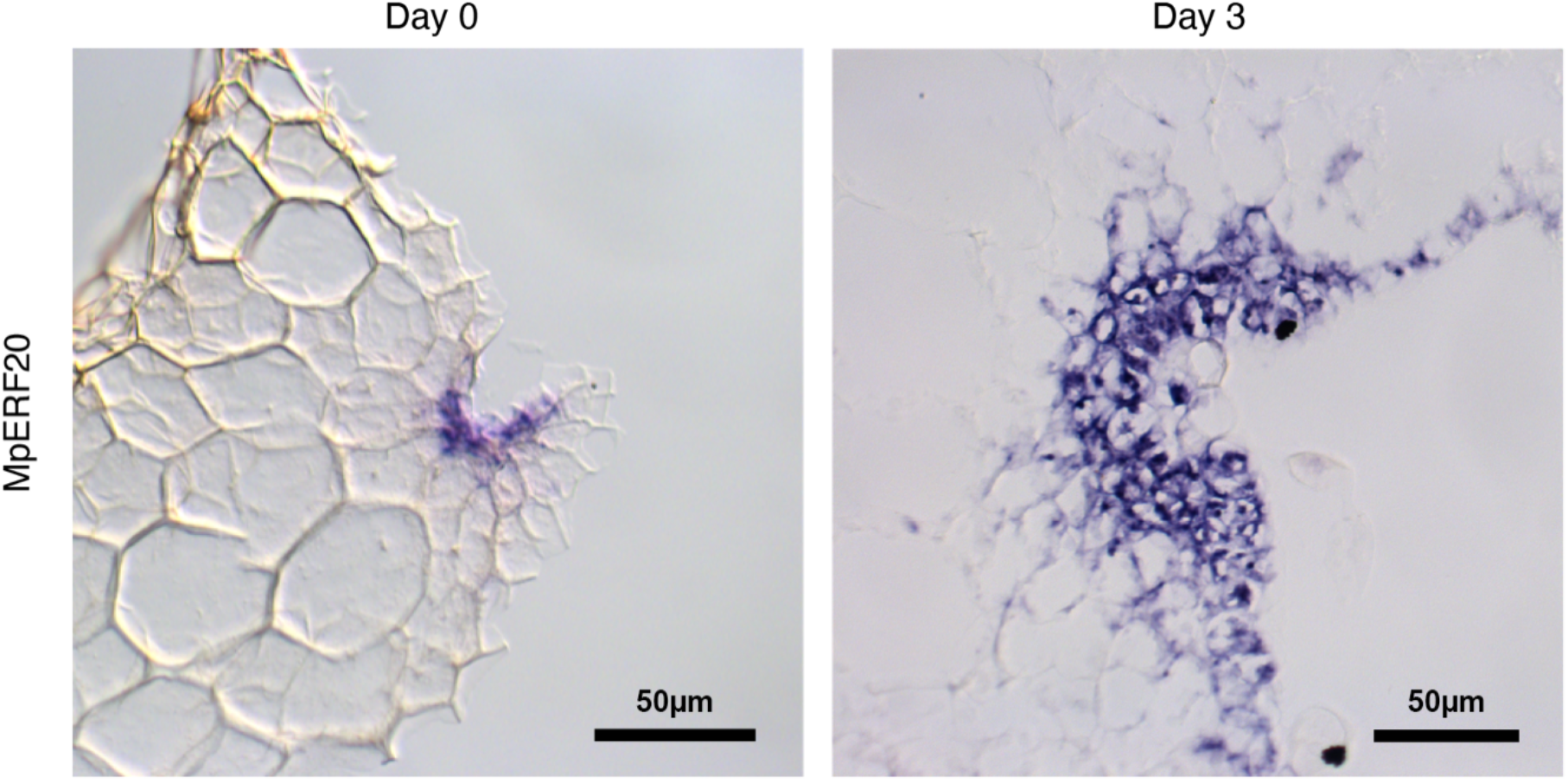
*In situ* localization of Mp*ERF20/LAXR* mRNA in 0-day-old and 3-day-old gemmalings. Signal is specifically localised in cells around the SCZ. Scale bar = 50 μm.

**Supplemental movie S1. Additional dynamic expression of reporters in the SCZ.** Time-lapse of *_pro_*Mp*ERF20/LAXR* (yellow) expression after laser ablation of the notches and until re-establishment of the new SCZ. Constitutive plasma membrane marker (*_pro_*Mp*UBE2:mScarlet-Lti6b*) is shown in magenta.

